# Nonlinear EEG signatures of mind wandering during breath focus meditation

**DOI:** 10.1101/2022.03.27.485924

**Authors:** Yiqing Lu, Julio Rodriguez-Larios

## Abstract

In meditation practices that involve focused attention to a specific object, novice practitioners often experience moments of distraction (i.e., mind wandering). Previous studies have investigated the neural correlates of mind wandering during meditation practice through Electroencephalography (EEG) using linear metrics (e.g., oscillatory power). However, their results are not fully consistent. Since the brain is known to be a chaotic/nonlinear system, it is possible that linear metrics cannot fully capture complex dynamics present in the EEG signal. In this study, we assess whether nonlinear EEG signatures can be used to characterize mind wandering during breath focus meditation in novice practitioners. For that purpose, we adopted an experience sampling paradigm in which 25 participants were iteratively interrupted during meditation practice to report whether they were focusing on the breath or thinking about something else. We compared the complexity of EEG signals during mind wandering and breath focus states using three different algorithms: Higuchi’s fractal dimension (HFD), Lempel-Ziv complexity (LZC), and Sample entropy (SampEn). Our results showed that EEG complexity was generally reduced during mind wandering relative to breath focus states. We conclude that EEG complexity metrics are appropriate to disentangle mind wandering from breath focus states in novice meditation practitioners, and therefore, they could be used in future EEG neurofeedback protocols to facilitate meditation practice.

## Introduction

In the last years, several studies have shown that different meditation practices can promote mental and physical health (Okoro et al., 2013; Tang et al., 2015; Khoury et al., 2017; Deolindo et al., 2020). In an important part of these meditation practices, practitioners focus their attention on a specific object of meditation (such as the sensation of breathing) and try to disengage from distractions (Lutz et al., 2015; Anālayo, 2019). These moments of distraction are usually categorized as ‘mind wandering’ and they entail self-generated thoughts about the past, present, and future (Smallwood and Schooler, 2006, 2015; Delorme and Brandmeyer, 2019). Mind wandering during meditation is more prominent in novice practitioners, who normally experience more difficulties to detect mind wandering episodes and to re-focus their attention on the object of meditation (Mrazek et al., 2013; Brandmeyer and Delorme, 2018; Linares Gutiérrez et al., 2019; Rodriguez-Larios et al., 2021).

There is a growing interest in the identification of the neural correlates of mind wandering during meditation through Electroencephalography (EEG). This is partly due to its potential to develop EEG-assisted meditation protocols (Brandmeyer and Delorme, 2013; Ros et al., 2013; Brandmeyer and Delorme, 2020). Several studies have adopted experience sampling paradigms to study the EEG correlates of mind wandering in the context of breath focus meditation (Braboszcz and Delorme, 2011; Brandmeyer and Delorme, 2018; van Son et al., 2019; Rodriguez-Larios and Alaerts, 2021; Rodriguez-Larios et al., 2021). For experience sampling during meditation, also see a recent systematic review by Wahbeh et al. (2018). In this type of paradigm, subjects are interrupted several times during meditation so they can report whether they were focusing on the object of meditation or thinking about something else. The majority of these studies have focused on linear EEG metrics such as power. In this way, although changes in the alpha (8-14 Hz) and theta (4-8 Hz) bands have been associated with mind wandering, the direction of the effects is not fully consistent (Braboszcz and Delorme, 2011; Brandmeyer and Delorme, 2018; van Son et al., 2019; Rodriguez-Larios and Alaerts, 2021; Rodriguez-Larios et al., 2021).

Although the neural correlates of different cognitive states are normally studied through linear metrics (e.g., oscillatory power), the brain is known to be a chaotic system that presents nonlinear dynamics (Zhang et al., 2001; Rodriguez-Bermudez and Garcia-Laencina, 2015). Therefore, nonlinear EEG metrics are appropriate to provide complementary information of neural dynamics and their underlying mechanisms beyond conventional spectral analysis (Pereda et al., 2005; Rodriguez-Bermudez and Garcia-Laencina, 2015; Ma et al., 2018). Nonlinear methods can be used to estimate the EEG complexity or entropy, which can be interpreted as the degree of randomness in brain activity. The application of nonlinear methods to EEG analysis has generated considerable interest in recent years given their ability to characterize both healthy and pathological brain activity (Gomez et al., 2009; Ibáñez-Molina and Iglesias-Parro, 2014; Hou et al., 2021).

The neurobiological and phenomenological correlates of EEG complexity are still debated. From a biological standpoint, EEG complexity is thought to be highly influenced by the number of EEG generators and the level of oscillatory synchronization (Ibáñez-Molina & Iglesias-Parro, 2014; Schaworonkow & Nikulin, 2022). In this way, if the EEG signal is dominated by a single rhythm across the cortex, complexity would be minimized. Recent literature also suggests that EEG complexity could be affected by the excitation:inhibition ratio in the brain (as reflected in the EEG power law exponent) (Medel et al., 2020). In this view, greater inhibition would be associated with a more pronounced 1/f slope of the EEG spectrum and lower complexity (Gao et al., 2017). From a phenomenological perspective, it has been proposed that the level of complexity in brain activity is positively associated with the vividness of subjective experience (Carhart-Harris et al., 2014; Carhart-Harris & Friston, 2019). This theory is based on research with psychedelic drugs, which have indeed shown to transiently increase EEG complexity (Timmermann et al., 2019).

To our knowledge, no previous study has investigated the relationship between EEG complexity and mind wandering in the context of meditation practice with novice meditators. Given previous inconsistencies with linear metrics (Braboszcz and Delorme, 2011; Brandmeyer and Delorme, 2018; van Son et al., 2019; Rodriguez-Larios and Alaerts, 2021; Rodriguez-Larios et al., 2021), studying the non-linear EEG correlates of mind wandering during meditation practice holds high promise from a translational perspective. In this way, EEG complexity could be an alternative to EEG linear metrics to develop EEG-neurofeedback protocols aimed at facilitating meditation practice in novice meditation practitioners (Brandmeyer & Delorme, 2013, 2020).

In this study, we used an experience sampling paradigm to investigate nonlinear EEG features of mind wandering during breath focus meditation in participants without previous meditation experience. Participants (N = 25) were asked to focus on the sensation of breathing and to report whether they were focusing on their breath or thinking about something else after they heard a bell sound. We compared EEG complexity between mind wandering (MW) and breath focus (BF) states. For this purpose, we adopted three different algorithms to calculate complexity: Higuchi’s fractal dimension (HFD), Lempel-Ziv complexity (LZC), and Sample entropy (SampEn) (Lempel and Ziv, 1976; Higuchi, 1988; Richman and Moorman, 2000). Our choice of complexity metrics is due to several factors. First, these three metrics are widely used in the EEG literature and therefore, it would facilitate comparing our results with previous findings. Second, these metrics can be applied to relatively short time series, which is normally the case in EEG studies that adopt experience sampling paradigms. Lastly, HFD, LCZ and SamEn have a relatively low computational cost, which could eventually facilitate their application in real-time EEG neurofeedback protocols.

## Methods

### Dataset

In this study, we further analyzed a publicly available EEG dataset uploaded in the OSF platform: https://osf.io/b6rn9/. For a detailed description of the methods see Rodriguez-Larios & Alaerts (2021).

### Participants

Twenty-eight participants (age 23.46 y, range 20-29 y, 11 males) took part in the study. All participants did not have previous experience with meditation techniques. Twenty-five participants were included for further analysis since three participants were rejected due to technical problems in the experiment. All participants gave written informed consent before the experiment, which was approved by the Social and Societal Ethics Committee (SMEC) of the University of Leuven and was conducted in accordance with the Declaration of Helsinki. Participants were paid 8 euros per hour.

### Task

All participants performed a breath focus meditation (with eyes closed) while EEG was recorded. The meditation was randomly interrupted with a bell sound (after varying intervals of 20 to 60 seconds). Participants were then required to open their eyes and report whether they were mind wandering (MW) or focusing on their breath (breath focus; BF). Two additional questions allowed participants to report their confidence in their answer about their current state and arousal level in each trial. Each participant performed a total of 40 trials (the experiment lasted for approximately 40 minutes). Questions were displayed on a computer screen and answers were given by key pressing (using E-prime 2.0 software). Debriefings for the level of drowsiness, arousal, and emotional valence were filled after the task (available for 22 out of 25 participants).

### EEG acquisition and pre-processing

The EEG was recorded using a Nexus-32 system (Mind Media) (21 EEG electrodes including two mastoids). EEG data were sampled at 512 Hz. Pre-processing was performed using custom scripts and the EEGLAB toolbox (Delorme and Makeig, 2004) in MATLAB. The data was bandpass filtered between 1 Hz and 40 Hz (function pop_eegfiltnew). The filter order is estimated automatically through a heuristic in the EEGLAB function ‘pop_eegfiltnew.m’. In our analysis, the resulting filter order was 1690 for the high pass filter and 170 for the low pass filter. Abrupt artifacts were corrected using the Artifact Subspace Reconstruction method (function clean_asr with a cut-off value of 20 SD; see Chang et al., 2020). Independent Component Analysis (ICA) was performed to correct for eye movements (Delorme and Makeig, 2004). Components were rejected based on their spatial topography and their correlation with H/VEOG electrodes. ICA led to an average removal of 1.68 ± SD 0.47 components per subject. Data were average-referenced and epochs (i.e., BF or MW) of 5 seconds before bell sound probes were selected for further analysis. Epochs with absolute amplitudes exceeding 100 μV were rejected.

### Estimation of EEG complexity

#### Higuchi’s fractal dimension (HFD)

Several algorithms of fractal dimension have been developed. In comparison with other algorithms, HFD (Higuchi, 1988) is a prominent method that is suitable for analyzing EEG signals, which could be deterministic, stochastic, non-stationary, and noisy (Klonowski, 2009). This is due to the following advantages: relatively higher accuracy, faster computation, and applicability with relatively shorter time series of EEG (Accardo et al., 1997; Esteller et al., 2001; Gomez et al., 2009; Ibáñez-Molina and Iglesias-Parro, 2014).

The original time series *X* with *N* points can be used to construct *k* time series as follows:

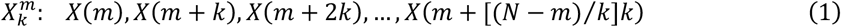

where *m* denotes the initial time (*m* = 1, 2, 3, …, *k*) and *k* denotes delay between the points. The length *L*_*m*_(*k*) is defined by

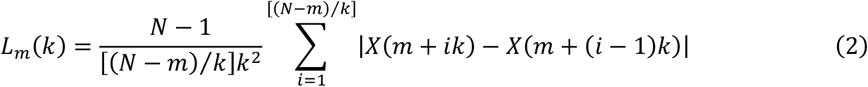

The average length *L*_*m*_(*k*) for *m* ranging from 1 to *k* is then obtained

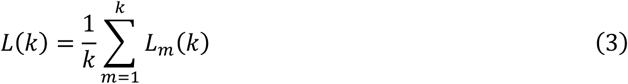

The *L*(*k*) is proportional to *k* ^*-D*^,

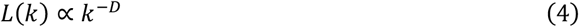

where *D* is the slope of the least squares linear fit through the curve of log*L*(*k*) versus log(1/*k*). The *D* is defined as the Higuchi fractal dimension HFD.

Note that *m* = 1, 2, 3, …, *k*, thus the *k* value must be determined, which is used in the analysis is defined as *k*_*max*_. To determine the *k*_*max*_, a range of *k*_*max*_ was applied to calculate the HFD, then a saturated HFD plotting indicated an appropriate *k*_*max*_. In the present study, we chose the *k*_*max*_ value of 80.

#### Lempel-Ziv complexity (LZC)

LZC was proposed by Lempel and Ziv (1976). LZC does not take into account whether the signal is from deterministic chaos or stochastic processes, i.e., model-independence, therefore it is appropriate for EEG signals (Zhang et al., 2001) and especially for the present study in which we do not aim to test for a model form of deterministic chaos or stochastic processes for our data, but rather interested in the distinct EEG complexity patterns of a specific experimental condition. Furthermore, the core part of the LZC algorithm just comprises sequence comparison and number accumulation. This can remarkably lower the requirement for computation cost, and eventually facilitates the application of this method, especially for real-time neurofeedback usage.

LZC algorithm depends on the process of coarse-graining by using which a data series is converted into a finite symbol sequence before calculating the complexity. We used binary conversion (“0–1”) in this study. The LZC can be estimated using the following steps: First, a time series with a length of *n* is binarised. The threshold can be the mean value or median value of the time series. Here we used the median value (of each channel of each epoch): *md*. The values of time series greater than *md* were assigned ones, lower than *md* were assigned zeros. Second, the binary sequence was scanned from left to right for different subsequences of consecutive characters which compose a vocabulary to summarise the series. The amount of different subsequences in the vocabulary is defined as the complexity counter *c*(*n*). Imagine a regular and simple signal that be binarised to a sequence of 010101010101, which can also be shown as 0*1*0*1*0*1*0*1*0*1*0*1 (subsequences separated with asterisks). Different subsequences are “0” and “1”, thus the complexity of this signal is *c*(*n*) = 2. Consider another example: an irregular signal be binarised to a more complex sequence such as 110100010110, which can be shown as 1*10*100*0101*10 (subsequences separated with asterisks). This signal has a higher number of different subsequences: “1”, “10”, “100”, and “0101”, reflecting that the complexity is relatively higher: *c*(*n*) = 4. Therefore, the basic idea of this algorithm just includes character comparison and number accumulation (Zhang et al., 2001).

To eliminate the effects of signal length, *c*(*n*) can be normalized by using the upper bound of complexity:

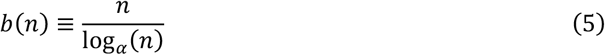

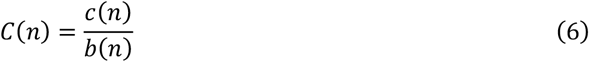

For binary conversion, *α* = 2. Here, *C*(*n*) is the LZC measure used in the present study.

#### Sample Entropy (SampEn)

Entropy is a concept from information theory, can measure the uncertainty of information/signal, can quantify the complexity of a system. Many algorithms have been developed and/or improved to estimate the entropy of a system. For biological time series data, approximate entropy (ApEn) (Pincus, 1991) and sample entropy (Richman and Moorman, 2000) are the two of the most commonly used measures (Yentes et al., 2013). There are several advantages of SampEn as compared to ApEn including better relative consistency for results, less sensitivity to the length of time series, remarkably faster computation, and without a bias towards regularity (Pincus, 1995; Richman and Moorman, 2000; Ramdani et al., 2009; Zhang et al., 2009; Yentes et al., 2013; Delgado-Bonal and Marshak, 2019).

The algorithm of SampEn is independent of the series length and can be described as follows. For a time series with *N* points *u*, a template vector of length *m* is defined:

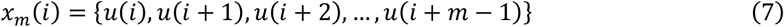

in which *i* = 1, 2, *N*-*m*+1. Then, we define:

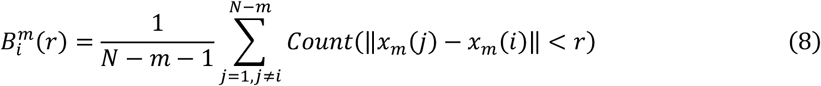

The sum is the number of vectors *x*_*m*_(*j*) within a distance *r* of *x*_*m*_(*i*). Here we used the Chebyshev distance. In order to exclude self-matches, set *j* ≠ *i*. The parameter *r* is the tolerance value for accepting matches, normally set as a portion of standard deviation (*σ*) of data. We next define the function *B*^*m*^(*r*):

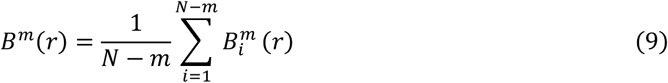

*B*^*m*^(*r*) is the probability that two vectors match for *m* points. Similarly, we applied the definitions for *m* + 1 points:

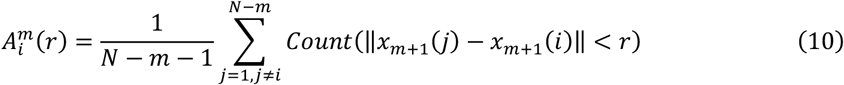

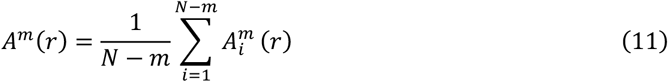

The *SampEn* is then defined as:

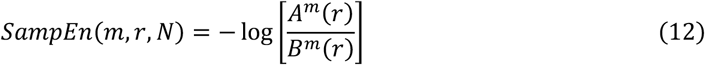

In the present study, we used *m* = 2, *r* = 0.2*σ*. The higher value of SampEn, the higher irregularity and complexity of the data series.

### Statistical analysis

We analyzed the EEG data in the time window involving the last 5 seconds before the bell sound probes, using custom scripts in MATLAB (R2018a). The values of HFD, LZC and SampEn were analyzed separately. For each participant, we calculated the weighted average values across epochs (weighted by the confidence level of each trial) within the same condition for each electrode. These values were then used to evaluate the differences between MW and BF conditions. Dependent samples *t*-test (two-tailed) were computed to evaluate condition-related differences. In order to resolve the multiple comparisons problem, the cluster-based non-parametric randomization test (Maris and Oostenveld, 2007) implemented in the MATLAB toolbox FieldTrip (Oostenveld et al., 2011) was used. All paired samples were computed and a *p*-value of 0.05 was set as the threshold. The samples with a *p*-value under the threshold were clustered based on spatial-temporal adjacency, then the sum of the clustered *t*-values was obtained. This step was then repeated 1000 times by using samples randomized across MW and BF conditions, generating a new distribution of statistical values. The original samples were evaluated based on this distribution by using the threshold of 0.05 for significant data cluster(s). In FieldTrip, the cluster-level statistic at the threshold of 0.05 (two-sided test) is by setting the parameters cfg.alpha = 0.025 and cfg.tail = 0. In addition, to resolve the multiple comparisons problem in correlation analysis, the false discovery rate (FDR) method was applied, according to Benjamini and Hochberg (1995). Data were plotted using FieldTrip and custom scripts.

Unequal trial counts in different conditions may introduce bias for analysis. To overcome this potential problem, we also matched trial counts (MTC) between MW and BF conditions for each subject by randomly discarding trials from the condition with a larger trial count and then performed a comparison of complexity measures for the count-matched trials between conditions as well. Six participants were rejected due to insufficient (<10) trial counts in both conditions after MTC. Data from the remaining nineteen participants were used for MTC processed analysis. Note that, due to a technical problem only sixteen of these participants had debriefings for the level of drowsiness at the end of the experiment. We called the results without MTC processing “non-MTC” for simplicity.

### Correlation analysis

Given the relatively high levels of drowsiness of some participants in the sample (see Rodriguez-Larios and Alaerts, 2021), we assessed whether condition effects in complexity measures were associated with inter-individual differences in self-reported drowsiness. For this purpose, we calculated the averaged difference (MW minus BF) of complexity measures in the identified significant clusters for each subject and performed Kendall’s correlations between the averaged difference and the level of drowsiness across subjects. In order to uncover the information hidden by significant clusters, we also assessed the correlations between differences in complexity measures and drowsiness levels, per electrode (the complexity value of each electrode) and for all electrodes (the mean value across all electrodes) across subjects.

Assuming that drowsiness increases over time, Spearman’s correlation was analyzed between trial complexity and trial serial number per subject, and the *ρ(rho)*-value was obtained for each subject (N = 25). Then a one-sample *t*-test (two-tailed) was performed over the *ρ(rho)*-values. If the H0 hypothesis cannot be rejected (at the 5% significance level), the complexity value is not affected by the trials over time.

To investigate whether condition effects in complexity measures were associated with the differences in the low-frequency (4-12 Hz) range, we also calculated the averaged difference (MW minus BF) in amplitude (both absolute amplitude and relative amplitude) in theta (4–7 Hz) and alpha (8–12 Hz) frequency range in identified significant clusters for each subject (N = 25, data from Rodriguez-Larios and Alaerts, 2021). Pearson’s correlations were then performed between the averaged difference of complexity measures and amplitudes across subjects.

## Results

### Higuchi’s fractal dimension (HFD)

We observed decreased HFD during MW compared to BF. The cluster statistics revealed that the decrease in complexity (*t*(24) = −52.442, *p* < 0.001) was widely distributed across electrodes (see Figure 2). For this cluster, the mean HFD value was 1.677 (std: 0.039, median: 1.675) for MW and 1.695 (std: 0.026, median: 1.693) for BF. The values for each subject are shown in the scatter plot of Figure 2.

**Figure 1.**
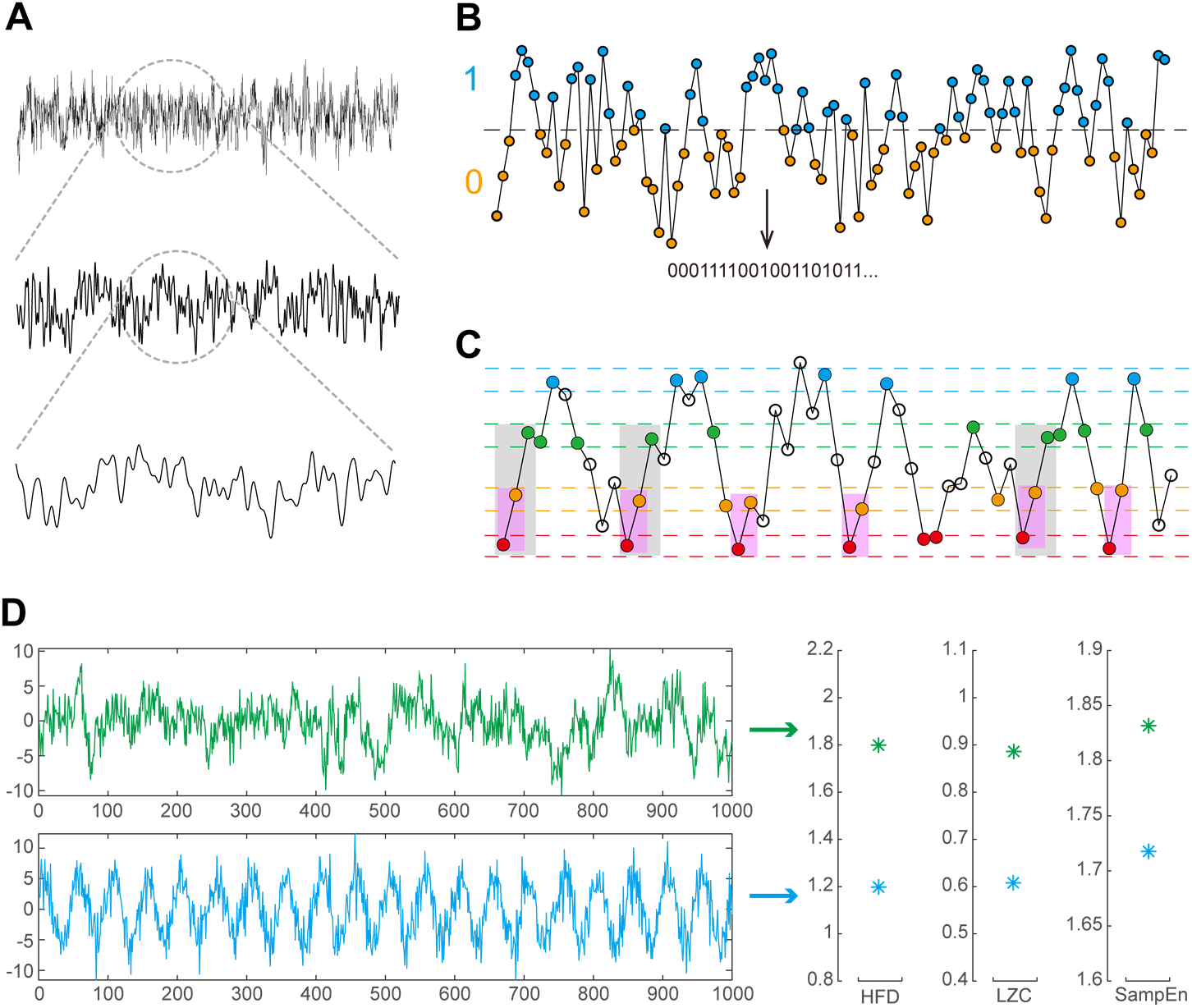
Illustration of the HFD, LZC and SampEn analysis. **(A)** The EEG time series reflect, to a certain degree, self-similarity, revealed at different timescales. The HFD is used to determine the fractal dimension of such data series. **(B)** In LZC analysis, a time series is binarised (by using the median value of each channel of each epoch). The values greater than the median are assigned ones (blue dots), and lower than the median are assigned zeros (orange dots). LZC can be defined as the number of unique subsequences in the binary sequence. This number can be normalized to eliminate the effects of signal length. This normalized value is the LZC used in this study. This panel was inspired by Leemburg and Bassetti (2018). **(C)** SampEn (here *m* = 2). Datapoints between dashed lines of the same color (red, orange, green and blue, respectively.) are denoted for accepting matches (i.e., the tolerance: 20% of standard deviation). Matching points are indicated by the corresponding color. Starting from the first data point, the number of the same sequence patterns of consecutive data points are counted, also highlighted by color bars: 2-consecutive datapoints (6 pink bars): red -> orange; 3-consecutive datapoints (3 gray bars): red -> orange -> green. This procedure is repeated from the second data point, and so on. Then the total number of 2- and 3-consecutive datapoint sequence patterns are determined respectively, and the natural logarithm of their ratio is SampEn. Intuitively, the more non-repeated patterns, the larger the SampEn. This panel was inspired by Costa et al. (2005). **(D)** Snippets of simulated signals (1000 data points each): a more complex signal (green) and a more regular signal (blue). The latter shows a more pronounced oscillatory pattern. The HFD, LZC and SampEn values of these two signals are shown on the right (for HFD analysis of the simulated signals: we use the *k*_*max*_ value of 30). It can be noticed that the values of the blue signal are lower than the values of the green signal respectively, reflecting a lower complexity of the blue signal.

**Figure 2.**
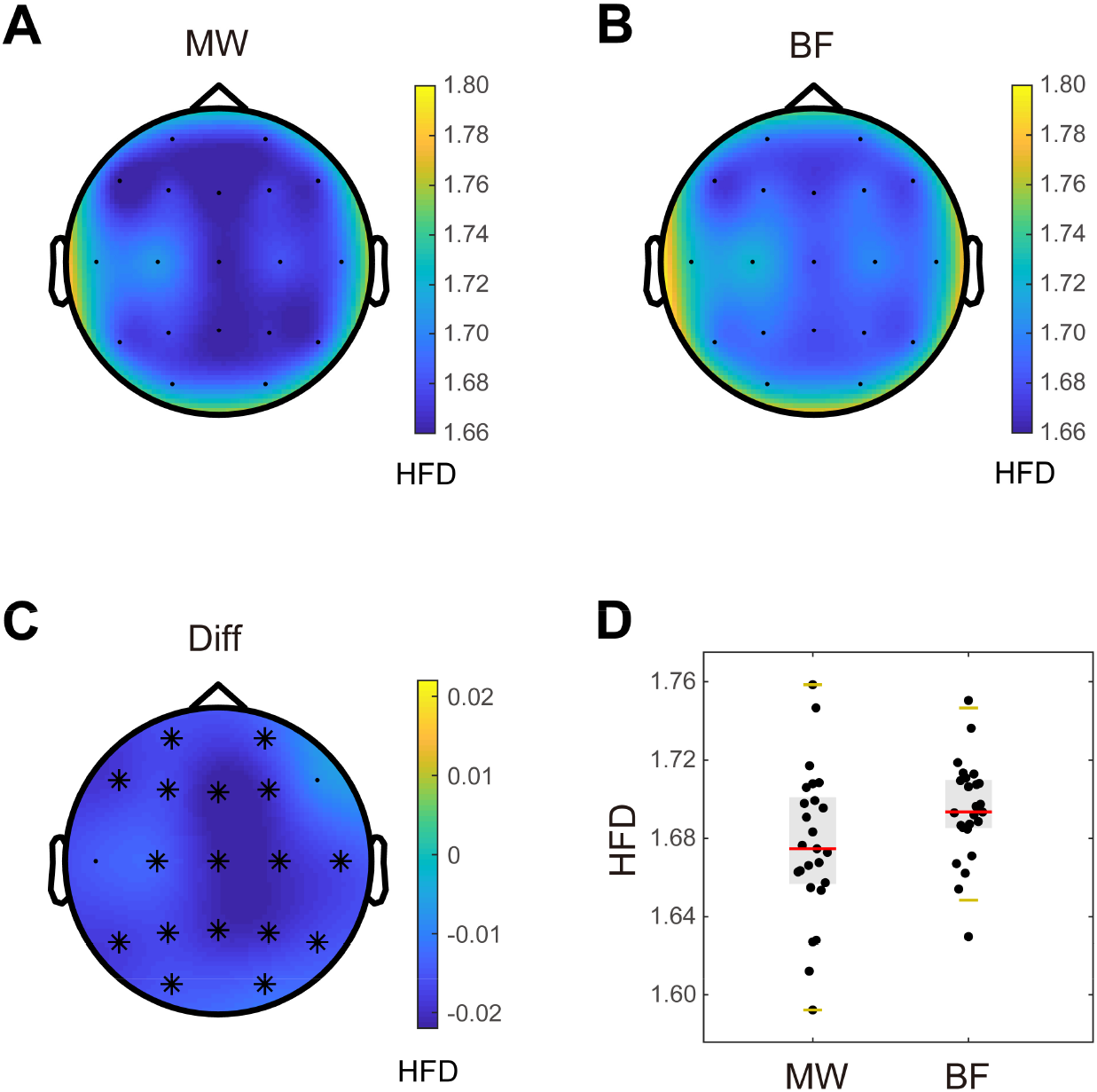
Differences in Higuchi’s fractal dimension (HFD) between mind wandering (MW) and breath focus (BF) conditions. **(A-B)** Topographical plots depicting mean HFD values for MW and BF conditions in each electrode. **(C)** Topographical plot depicting the mean difference (Diff) between MW and BF conditions in each electrode. The black asterisks mark electrodes in which the HFD decrease (MW < BF) was significant (*p* < 0.05). **(D)** Individual HFD values (averaged within the significant cluster) for MW and BF conditions. Each subject is represented by a dot. The gray boxes indicate the 25th and 75th percentiles. Centerlines show the median in each condition. When outliers are present, the whiskers indicate 1.5 times the interquartile range from the 25th and 75th percentiles. When no outliers are present, the whiskers lay on the most extreme data points.

Matched trial counts (MTC) analysis revealed a similar pattern of results. Decreased HFD (*t*(18) = −45.746, *p* = 0.004) was also observed during the MW condition relative to the BF condition across electrodes (Figure S1). The mean HFD value of the cluster was 1.674 for MW (std: 0.034, median: 1.671) and 1.696 (std: 0.018, median: 1.695) for BF. The values for each subject are also shown in Figure S1. This indicates that unequal trial counts did not affect the HFD results.

### Lempel-Ziv complexity (LZC)

A significant decrease in LZC was observed during MW relative to BF. Two negative clusters were found (MW < BF) (Figure 3). The first cluster revealed a significant (*t*(24) = −9.328, *p* = 0.020) LZC decrease (MW < BF) in the parietal and right frontocentral area (i.e., F4, Cz, C4, Pz). For this cluster, the mean LZC value for MW was 0.271 (std: 0.020, median: 0.273) while the mean LZC value for BF was 0.278 (std: 0.016, median: 0.279). The LZC values of this cluster for each subject are shown in the scatter plot of Figure 3. The second cluster (in the left frontal area) did not reach statistical significance (*t*(24) = −4.420, *p* = 0.057) (Fp1, F7).

**Figure 3.**
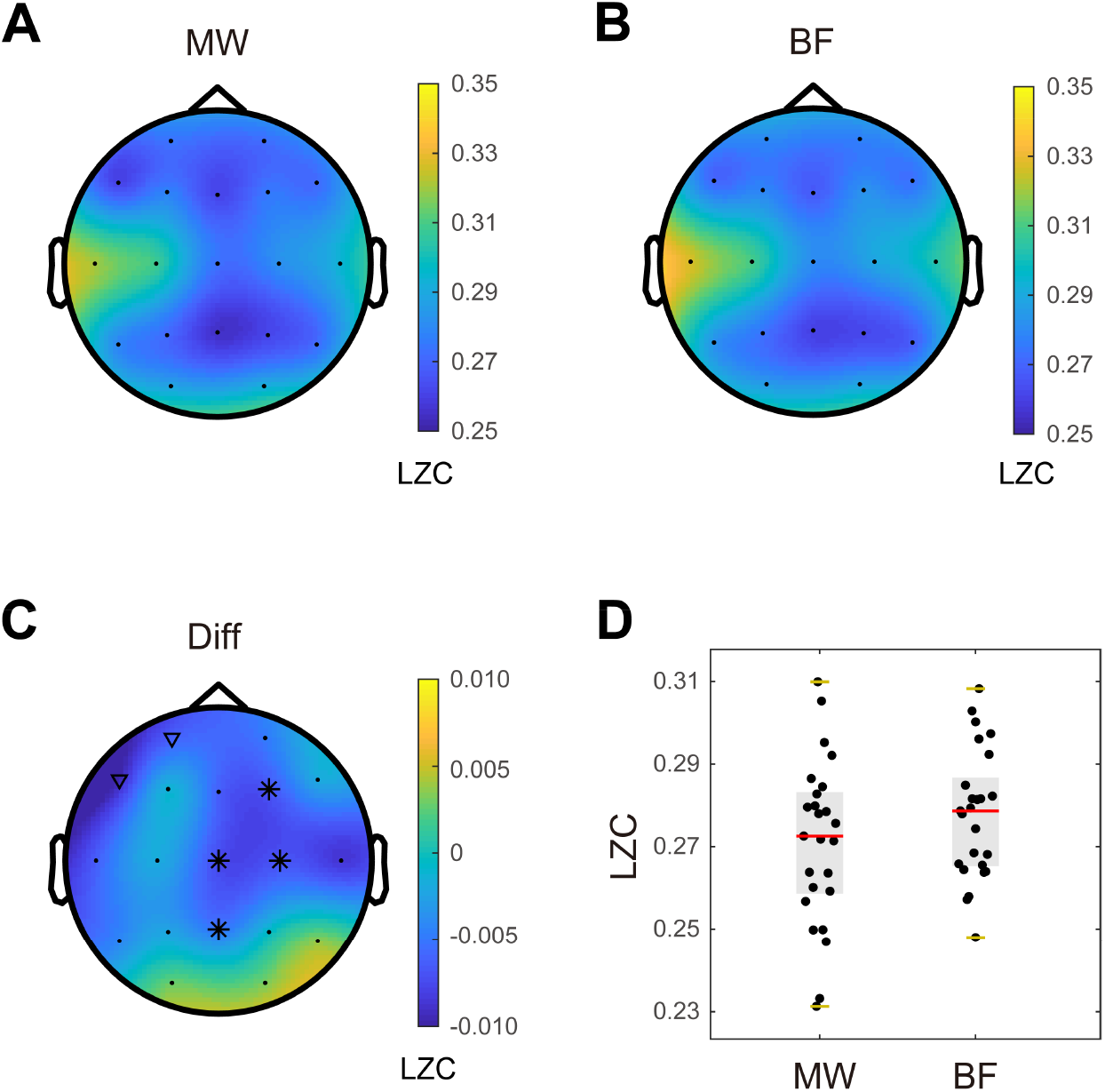
Differences in Lempel-Ziv complexity (LZC) between mind wandering (MW) and breath focus (BF) conditions. **(A-B)** Topographical plots depicting mean LZC values for MW and BF conditions in each electrode. **(C)** Topographical plot depicting the mean difference (Diff) between MW and BF conditions in each electrode. The black asterisks mark electrodes in which the LZC decrease (MW < BF) was significant (*p* < 0.05). The black triangles represent the second identified cluster which shows a not significant trend (*p* = 0.057) of LZC decrease (MW < BF) in the left frontal area. **(D)** Individual LZC values (averaged within the significant cluster, i.e., represented by black asterisks in Panel C) for MW and BF conditions. Each subject is represented by a dot. The gray boxes indicate the 25th and 75th percentiles. Centerlines show the median in each condition. When outliers are present, the whiskers indicate 1.5 times the interquartile range from the 25th and 75th percentiles. When no outliers are present, the whiskers lay on the most extreme data points.

When adopting the MTC approach, we also obtained two negative clusters (MW < BF) (Figure S2), although they did not reach statistical significance (*p* > 0.05). The first cluster showed a trend-level LZC decrease (*t*(18) = −4.248, *p* = 0.058) (MW<BF) in the parietal and right central area (i.e., C4, Pz). For this cluster, the mean LZC value was 0.267 for MW (std: 0.016, median: 0.267) and 0.273 for BF (std: 0.012, median: 0.272). Figure S2 shows the LZC values of this cluster for each subject. The second cluster showed a non-significant LCZ decrease (*t*(18) = −2.314, *p* = 0.154) (MW < BF) in the left frontal area (Fp1). In summary, although the LZC change became not significant (i.e., statistical trend) after controlling for trial counts, the direction and topography of LZC decreases (MW < BF) were qualitatively similar.

### Sample Entropy (SampEn)

A significant decrease in SampEn was observed during MW relative to BF (Figure 4). The cluster statistics revealed that the decrease (*t*(24) = −20.041, *p* = 0.007) occurred in frontal, right temporal, central, and parietal areas (i.e., Fp1, F7, Fz, F4, Cz, C4, T8, P4). The mean SampEn value of this cluster was 0.475 for MW (std: 0.031, median: 0.474) and 0.487 for BF (std: 0.024, median: 0.486) (see Figure 4).

**Figure 4.**
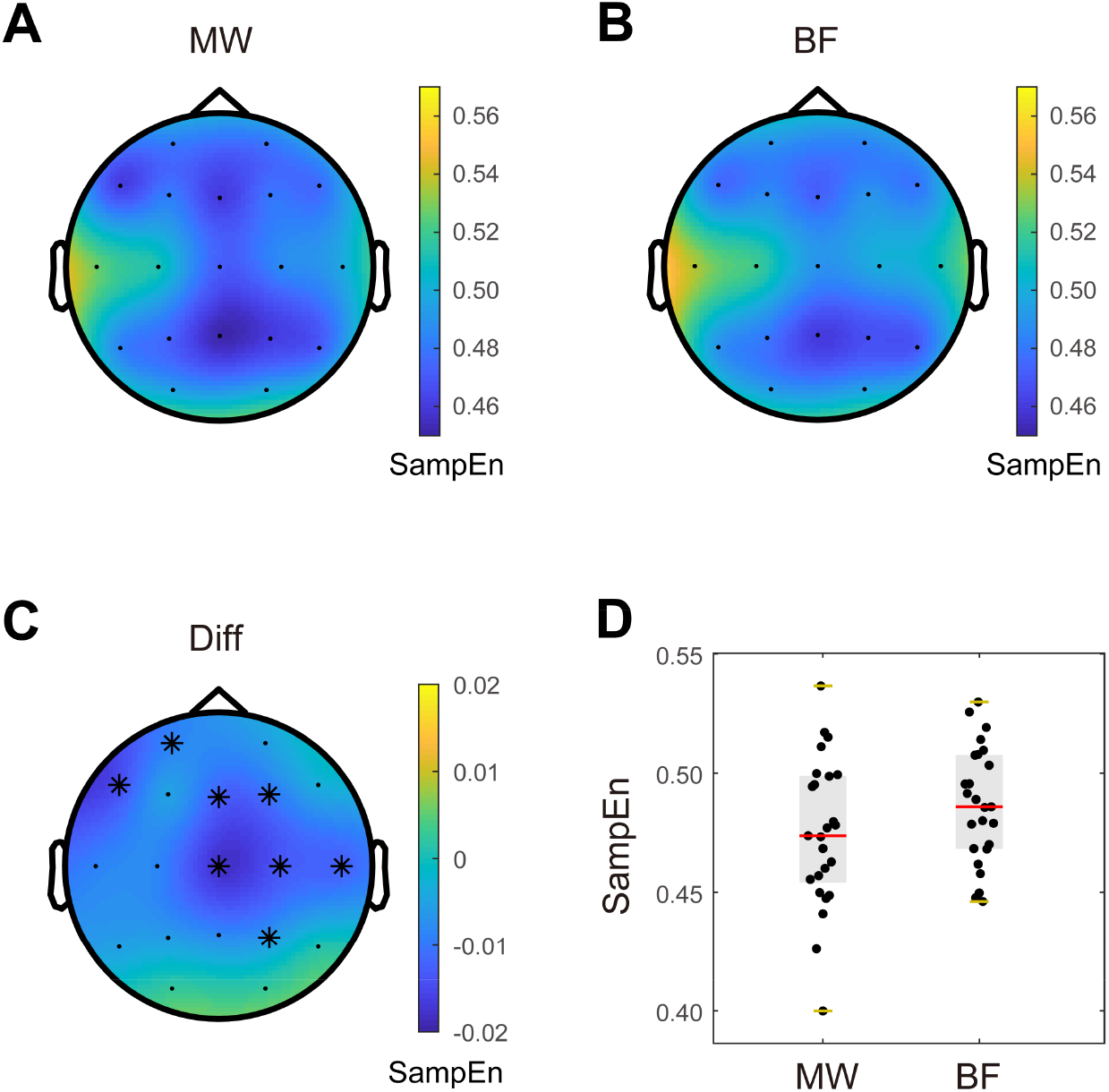
Differences in Sample entropy (SampEn) between mind wandering (MW) and breath focus (BF) conditions. **(A-B)** Topographical plots depicting mean SampEn values for MW and BF conditions in each electrode. **(C)** Topographical plot depicting the mean difference (Diff) between MW and BF conditions in each electrode. The black asterisks mark electrodes in which the SampEn decrease (MW < BF) was significant (*p* < 0.05). **(D)** Individual SampEn values (averaged within the significant cluster) for MW and BF conditions. Each subject is represented by a dot. The gray boxes indicate the 25th and 75th percentiles. Centerlines show the median in each condition. When outliers are present, the whiskers indicate 1.5 times the interquartile range from the 25th and 75th percentiles. When no outliers are present, the whiskers lay on the most extreme data points.

The MTC analysis revealed a significant decrease in SampEn (*t*(18) = −18.171, *p* = 0.009) during MW relative to BF in midline, right central and left frontal areas (i.e., Fp1, F7, Fz, Cz, C4, Pz) (Figure S3). The mean SampEn value in the cluster was 0.464 for MW (std: 0.028, median: 0.463) and 0.480 for BF (std: 0.020, median: 0.478). SampEn values for each subject are shown in Figure S3. In summary, similar to HFD, controlling for trial counts did not affect SampEn results (i.e., MW < BF).

### Relation between changes in complexity and drowsiness

In the light of previous results showing the effect of drowsiness in the EEG correlates of mind wandering (Rodriguez-Larios and Alaerts, 2021; Rodriguez-Larios et al., 2021), we also assessed how changes in complexity are associated with inter-individual differences in drowsiness levels. All three complexity measures showed a negative relationship with drowsiness (Figure S4), i.e., a greater decrease in complexity was associated with higher levels of drowsiness. Note that the correlation between complexity and drowsiness reached statistical significance for HFD (Kendall’s coefficient *τ* = −0.348, *p* = 0.036) but not for LZC or SampEn (*τ* = −0.177, *p* = 0.295 and *τ* = −0.205, *p* = 0.222 respectively). The previous results were derived from the significant clusters. Additionally, we found at the single-electrode level that C3, P3 and P4 showed a negative correlation (all *p* < 0.01) between the difference in HFD and the drowsiness (Table S1) after FDR correction. The correlation between complexity and drowsiness did not reach statistical significance for other single-electrode, nor the mean complexity values across all electrodes.

When adopting the MTC approach, we observed that all the complexity measures showed a negative relation to drowsiness, although in this case they did not reach statistical significance (for HFD, LZC and SampEn: *τ* = −0.293, *p* =0.153; *τ* = −0.248, *p* = 0.233 and *τ* = −0.218, *p* = 0.295, respectively) (Figure S4). Note that, since the LZC metric showed no significant clusters when comparing MW and BF conditions with the MTC approach, we used the cluster that presented a statistical tendency (*p* = 0.058; see Figure S2) to run the correlation with drowsiness. Similarly, for the MTC approach, the correlation analysis was also performed at the single-electrode level and the all-electrode level, and no significant result was found (Table S2).

If changes in complexity metrics reflect changes in drowsiness and/or fatigue, we could assume that complexity would decrease over time within subjects (as drowsiness would be expected to increase). However, we found no significant correlation (all *p* > 0.5) between EEG complexity (HDF, LZC, and SampEn) and trial numbers (Table S3). This result indicates that changes in complexity cannot be fully explained by drowsiness or fatigue.

## Discussion

In this study, we investigated nonlinear EEG features of mind wandering during breath focus meditation in participants without previous meditation experience. For this purpose, we adopted an experience sampling paradigm in which participants were repeatedly probed during a breath focus meditation to report whether they were focusing on their breath (BF) or thinking about something else (MW). Different EEG metrics (HFD, LZC and SampEn) revealed a significant decrease in complexity during MW relative to BF states. While differences in HFD were widespread across electrodes, effects in LZC and SampEn were more pronounced in central electrodes. In addition, our results also revealed that participants that reported higher levels of drowsiness tended to show a greater decrease in complexity during MW relative to BF states. However, these latter correlations rendered not significant when controlling for differences in the number of trials between conditions.

We here demonstrate that EEG activity is more predictable (i.e. less complex/random/entropic) during lapses of attention in the context of meditation practice. A more predictable EEG signal could be due to at least three different factors: i) reduced number of brain generators (Schaworonkow, & Nikulin, 2022) ii) increase in the power law exponent (Medel et al., 2020) and/or iii) greater presence of oscillatory activity (Timmermann et al., 2019). Concerning the number of generators, we can speculate that during mind wandering a specific network dominates the EEG signal and that is why complexity is reduced. Given its consistent association with mind wandering, a good candidate for this would be the Default Mode Network (DMN) (Brewer et al., 2011; Ellamil et al., 2016). On the other hand, if reduced EEG complexity is due to increases in the power law exponent and/or the presence of oscillatory activity (Medel et al., 2020), it is likely that this is reflecting increased cortical inhibition (Klimesch et al., 2007; Gao et al., 2017). In this line, lapses of attention have been previously associated with both low-frequency power increases and decreased excitability of the cortex (Braboszcz & Delorme, 2011; Smallwood et al.,2008). In this regard, it is important to note that our previous analysis of this data set indeed revealed a relative increase in low-frequency power during mind wandering relative to breath focus (which could be reflective of increased oscillatory activity and/or a more pronounced slope of the power law exponent) (Rodriguez-Larios & Alaerts, 2021). Our analysis has revealed that this increase is negatively correlated with reduced EEG complexity (Figure S5).

The ‘entropic brain’ theory posits that there is a correspondence between the ‘richness’ of brain activity and subjective experience (Carhart-Harris et al., 2014; Carhart-Harris & Friston, 2019). According to this theory, when brain activity is more diverse (higher entropy/complexity) subjective experience is more vivid. In support of this idea, it has been shown that after psychedelics intake both the EEG signal and subjective experience become more complex and disorganized (Timmermann et al., 2019). Given the similarities between meditative and psychedelic states (Millière et al., 2018), the entropic brain theory predicts that meditation should also increase complexity in brain activity (Carhart-Harris & Friston, 2019). In this line, we here show that moments of focused meditation in novice meditators have a relatively higher EEG complexity than moments of distraction. However, this is not fully consistent with previous literature with experienced meditators. Although some studies have indeed associated meditative states with higher EEG complexity (Kakumanu et al., 2018; Vivot et al., 2020), other studies have reported the opposite effect (Aftanas and Golocheikine, 2002; Young et al., 2021). It is possible that inconsistencies regarding the relationship between EEG complexity and meditative states are due to differences in the meditation tradition, the level of expertise and the adopted complexity metric (Aftanas and Golocheikine, 2002; Huang and Lo, 2009; Kakumanu et al., 2018; Kumar et al., 2020; Vivot et al., 2020; Young et al., 2021). Given the great number of possible EEG metrics/traditions/levels of expertise that can be assessed, the only way of achieving a consensus in this field would be to make raw EEG data from different studies publicly available thereby allowing to assess these factors systematically.

In addition to mind wandering, lapses of attention can occur because of drowsiness (Brandmeyer & Delorme, 2018). Crucially, decreases in complexity have also been reported during states of transition from wakefulness to sleep (Hou et al., 2021). Hence, it is possible that (at least part of) the self-reported mind wandering in our participants is due to drowsiness. We assess this possibility by correlating inter-individual differences in complexity changes (mind wandering – breath focus) and the level of drowsiness. Although we found that subjects with higher drowsiness tended to have a more pronounced reduction in complexity during mind wandering, this latter relationship rendered not significant when controlling for different trial counts between conditions. It is important to note that this latter correlational analysis was performed with a relatively small sample (N = 16, for matched trial counts) because drowsiness scores were not available in all subjects. Moreover, drowsiness was only reported at the end of the task, which did not allow us to assess whether EEG complexity and drowsiness covary within subjects throughout the task. Consequently, these results have to be interpreted with caution and further research is needed to disentangle mind wandering and drowsiness effects on complexity. Specifically, future studies using experience sampling could ask participants for their level of drowsiness (in addition to mind wandering) on a trial-by-trial basis. This would allow to assess the relationship between mind wandering and complexity while controlling for variations in drowsiness.

The identification of a reliable EEG correlate of attentional lapses during meditation could promote the development of EEG-neurofeedback protocols aimed at facilitating meditation practice (Brandmeyer and Delorme, 2013; Ros et al., 2013; Badran et al., 2017; Brandmeyer and Delorme, 2020). Since we find reduced EEG complexity during mind wandering relative to breath focus states in novices, complexity metrics seem adequate for this purpose. In this way, participants could be alerted of an attentional lapse through auditory and/or visual feedback if there is a relative reduction in EEG complexity (regardless this is due to mind wandering or drowsiness). This could be specially relevant for some clinical and non-clinical populations that show special difficulties to practice meditation due to their inability to control the occurrence of mind wandering (Zylowska et al., 2008; Cachia et al., 2016).

In conclusion, our results showed a significant decrease in EEG complexity during mind wandering relative to breath focus states in novice meditation practitioners. Based on previous literature, we speculate that increased predictability of the EEG signal during mind wandering (less randomness/complexity/entropy) could be due to a reduced number of cortical generators and/or increased overall cortical inhibition. From a translational perspective, our findings suggest that nonlinear EEG features (i.e., HFD, LZC, and SampEn) could be effectively used to facilitate meditation practice through EEG-neurofeedback.

## Abbreviations

ApEn: approximate entropy;
BF: breath focus;
Ctrl: control;
Diff: the difference between mind wandering and breath focus (MW minus BF);
DMN: Default Mode Network;
EEG: electroencephalography;
FDR: false discovery rate;
HFD: Higuchi’s fractal dimension;
ICA: independent component analysis;
LZC: Lempel-Ziv complexity;
MTC: matched trial counts;
MW: mind wandering;
SampEn: sample entropy;
SMEC: Social and Societal Ethics Committee.

## Acknowledgments

The authors would like to thank Hannah Bernhard for her helpful comments.

## Data and code availability statement

Raw EEG data can be found here: https://osf.io/b6rn9/. Codes used for this study are open to access at https://github.com/y-q-l/MW-BF-NL and https://osf.io/b6rn9/.

## Author contributions (CRediT)

**Yiqing Lu:** Conceptualization, Methodology, Investigation, Software, Formal analysis, Visualization, Validation, Writing – Original draft, Writing – Review & Editing. **Julio Rodriguez-Larios:** Conceptualization, Methodology, Software, Data curation, Resources, Writing – Review & Editing.

