## Supplementary Materials for "Nonlinear EEG signatures of mind wandering during breath focus meditation"

The supplementary materials include Figure S1-S5 and Table S1-S3.

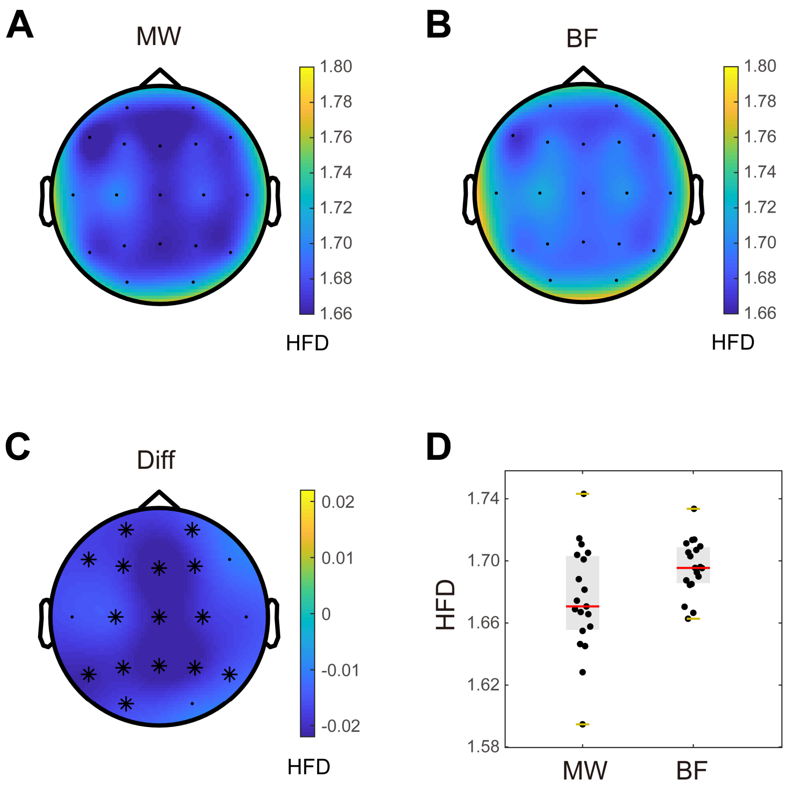


**Figure S1. Differences in Higuchi's fractal dimension (HFD) between mind wandering (MW) and breath focus (BF) conditions, when adopting the matched trial counts (MTC) approach. (A-B)** Topographical plots depicting mean HFD values for MW and BF conditions in each electrode. **(C)** Topographical plot depicting the mean difference (Diff) between MW and BF conditions in each electrode. The black asterisks mark electrodes in which the HFD decrease (MW < BF) was significant ($p < 0.05$). **(D)** Individual HFD values (averaged within the significant cluster) for MW and BF conditions. Each subject is represented by a dot. The gray boxes indicate the 25th and 75th percentiles. Centerlines show the median in each condition. When outliers are present, the whiskers indicate 1.5 times the interquartile range from the 25th and 75th percentiles. When no outliers are present, the whiskers lay on the most extreme data points.

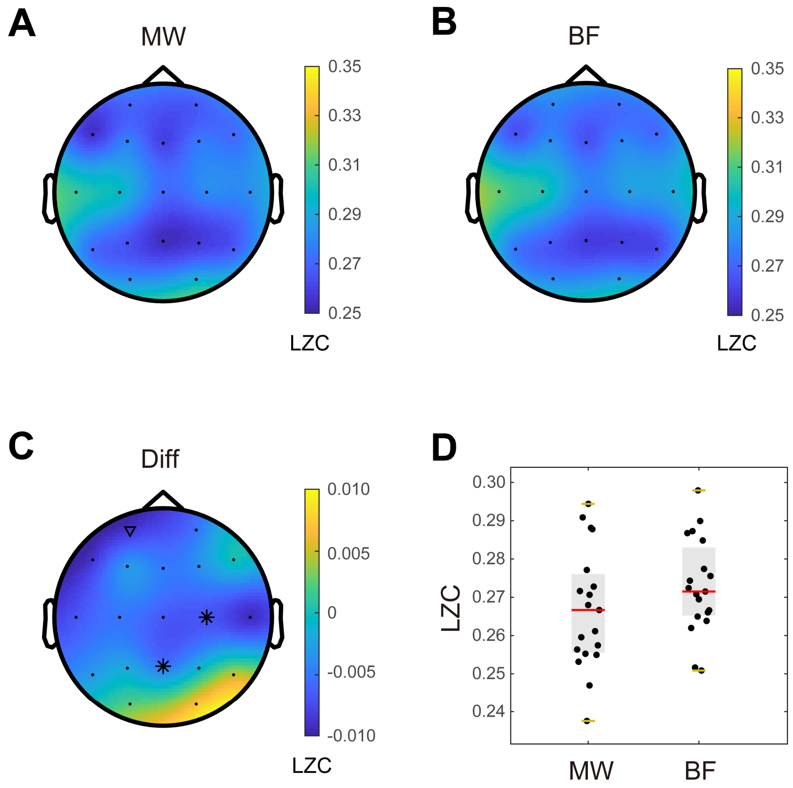


**Figure S2. Differences in Lempel-Ziv complexity (LZC) between mind wandering (MW) and breath focus (BF) conditions, when adopting the matched trial counts (MTC) approach. (A-B)** Topographical plots depicting mean LZC values for MW and BF conditions in each electrode. **(C)** Topographical plot depicting the mean difference (Diff) between MW and BF conditions in each electrode. The black asterisks and black triangle represent the first and the second identified clusters which show non-significant trends ($p$ = 0.058 and $p$ = 0.154, respectively) of LZC decrease (MW < BF). **(D)** Individual LZC values (averaged within the first identified cluster, i.e., represented by black asterisks in Panel C) for MW and BF conditions. Each subject is represented by a dot. The gray boxes indicate the 25th and 75th percentiles. Centerlines show the median in each condition. When outliers are present, the whiskers indicate 1.5 times the interquartile range from the 25th and 75th percentiles. When no outliers are present, the whiskers lay on the most extreme data points.

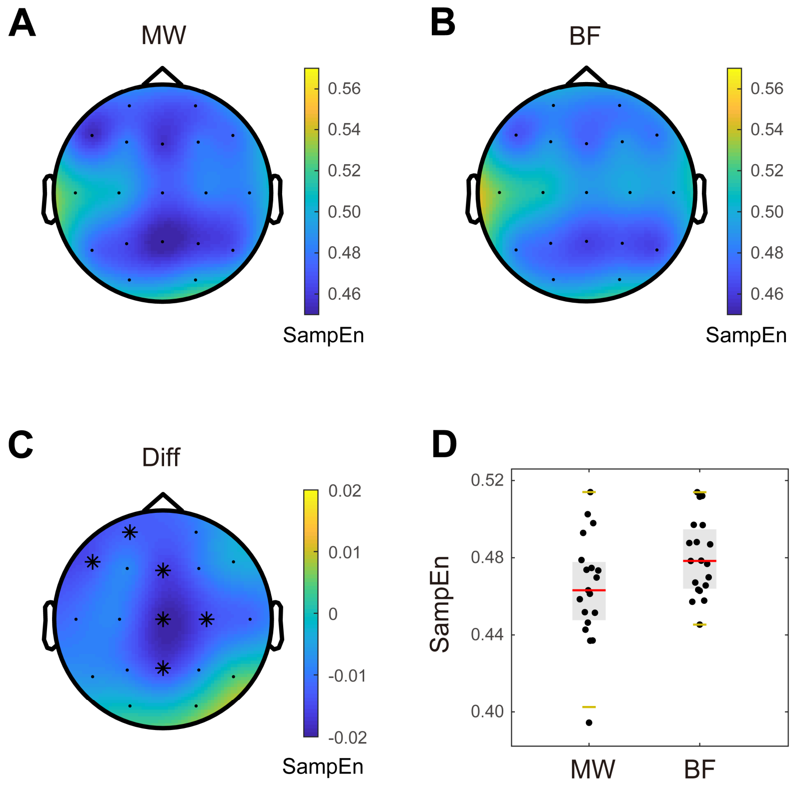


**Figure S3. Differences in Sample entropy (SampEn) between mind wandering (MW) and breath focus (BF) conditions, when adopting the matched trial counts (MTC) approach. (A-B)** Topographical plots depicting mean SampEn values for MW and BF conditions in each electrode. **(C)** Topographical plot depicting the mean difference (Diff) between MW and BF conditions in each electrode. The black asterisks mark electrodes in which the SampEn decrease (MW < BF) was significant ($p < 0.05$). **(D)** Individual SampEn values (averaged within the significant cluster) for MW and BF conditions. Each subject is represented by a dot. The gray boxes indicate the 25th and 75th percentiles. Centerlines show the median in each condition. When outliers are present, the whiskers indicate 1.5 times the interquartile range from the 25th and 75th percentiles. When no outliers are present, the whiskers lay on the most extreme data points.

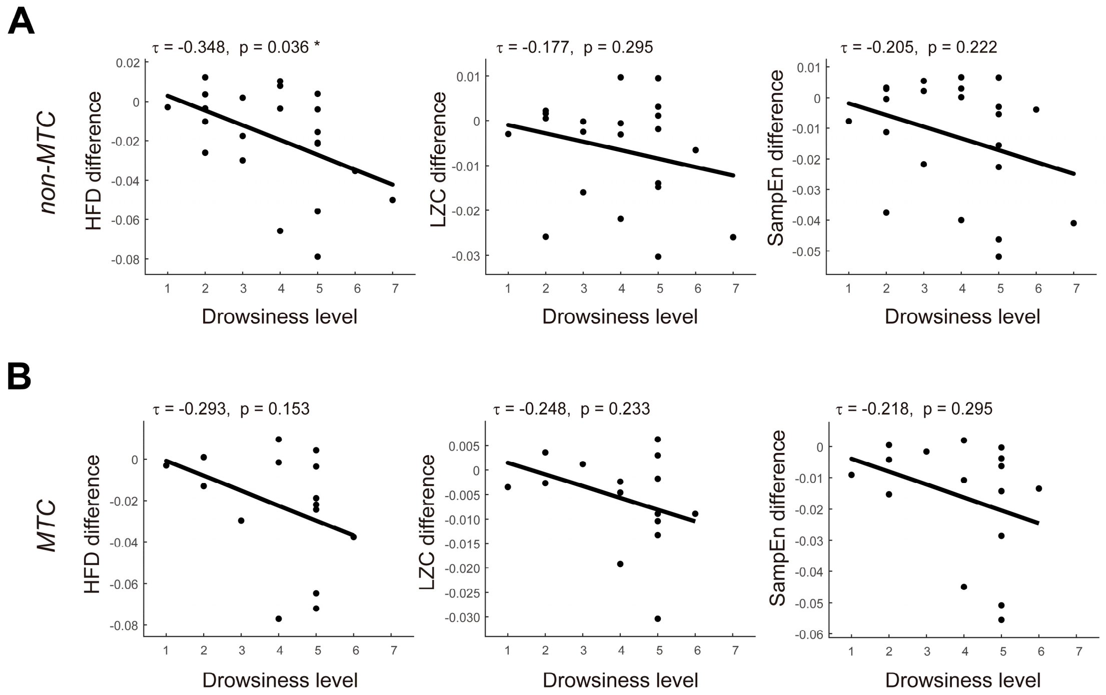


**Figure S4. Correlations between condition-related changes in EEG complexity and drowsiness level. (A-B)** non-MTC (non-matched trial counts) approach and MTC approach, respectively. In each panel, the difference in complexity (MW – BF) for each metric (HDF, LZC, and SampEn) is plotted as a function of drowsiness level. Each dot represents one subject (average complexity across electrodes showing significant condition effects). (For LZC, since the LZC metric showed no significant clusters when comparing MW and BF conditions with the MTC approach, the cluster that presented a statistical tendency ($p$ = 0.058) was used). Kendall's correlation coefficients and $p$-values are shown for each metric at the top of their corresponding panel. Although a negative correlation was observed for all metrics (i.e., a greater decrease in complexity during MW was associated with greater drowsiness), only HFD reached statistical significance ($p$ < 0.05 *) for the non-MTC approach. No statistical significance was reached for the MTC approach.

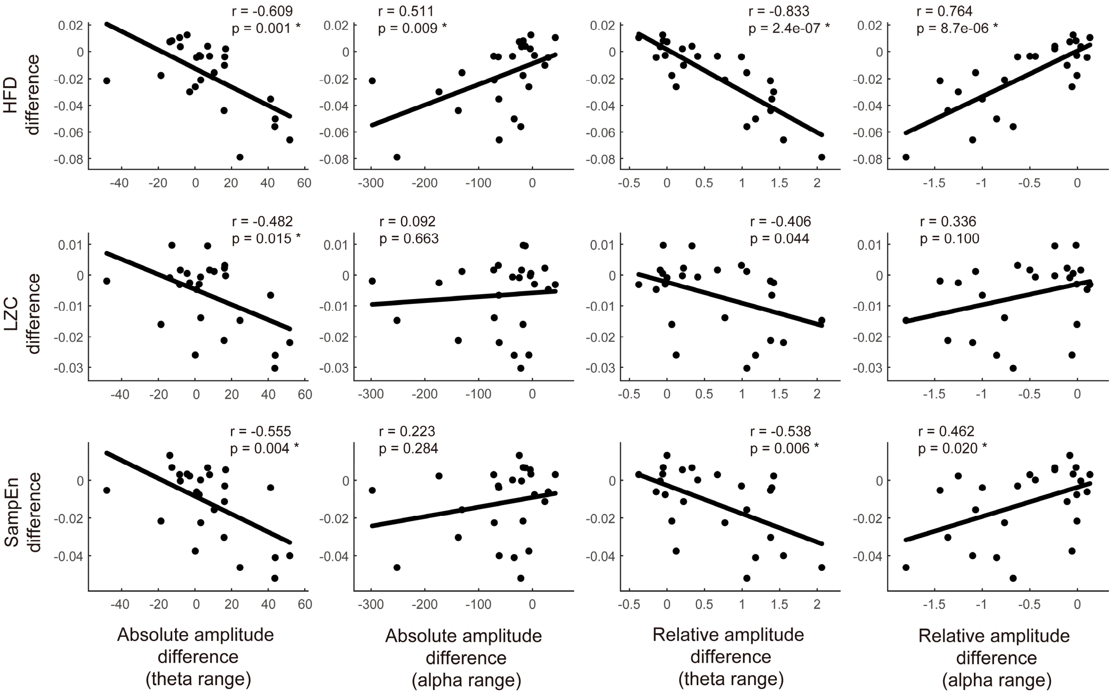


**Figure S5. Correlations between condition-related changes in EEG complexity and amplitude modulations in the theta-alpha frequency range.** Theta: 4–7 Hz; alpha: 8–12 Hz. In each panel, the difference (MW – BF) in complexity for each metric (HDF, LZC, and SampEn) is plotted as a function of the difference (MW – BF) in amplitude (absolute amplitude: columns 1 and 2; relative amplitude: columns 3 and 4). Each dot represents one subject (mean values across electrodes showing significant condition effects). The amplitude values are from our previous study (Rodriguez-Larios and Alaerts, 2021). Pearson's correlation coefficients and *p*-values (the original *p*-values, uncorrected by FDR) are shown at the top of their corresponding panels. Asterisks indicate the presence of significance after FDR correction (threshold: 0.05). A negative correlation was observed between complexity and amplitude modulations in the theta range, i.e., a decrease in complexity during MW was associated with an increase in theta power (columns 1 and 3). On the contrary, a positive correlation was observed between complexity and amplitude modulations in the alpha range, i.e., a decrease in complexity during MW was associated with a decrease in alpha power (columns 2 and 4). Note: a positive correlation was observed between LZC and alpha power (both for absolute amplitude and relative amplitude), but they did not reach statistical significance.

| Electrode | HFD | | LZC | | SampEn | |
|---|---|---|---|---|---|---|
| | *τ* | *p* | *τ* | *p* | *τ* | *p* |
| Fp1 | -0.243 | 0.146 | 0.109 | 0.523 | 0.109 | 0.523 |
| Fp2 | -0.233 | 0.163 | 0.071 | 0.684 | -0.005 | 1.000 |
| F7 | -0.119 | 0.486 | 0.005 | 1.000 | -0.005 | 1.000 |
| F3 | -0.233 | 0.163 | -0.043 | 0.816 | 0.052 | 0.771 |
| Fz | -0.395 | 0.017 | -0.109 | 0.523 | -0.186 | 0.270 |
| F4 | -0.195 | 0.245 | -0.052 | 0.771 | 0.014 | 0.954 |
| F8 | -0.119 | 0.486 | 0.100 | 0.561 | -0.148 | 0.383 |
| T7 | -0.205 | 0.222 | -0.129 | 0.450 | -0.109 | 0.523 |
| C3 | -0.452 | 0.006 * | -0.252 | 0.131 | -0.357 | 0.032 |
| Cz | -0.167 | 0.323 | -0.167 | 0.323 | -0.233 | 0.163 |
| C4 | -0.243 | 0.146 | -0.176 | 0.296 | -0.328 | 0.048 |
| T8 | -0.243 | 0.146 | -0.262 | 0.117 | -0.252 | 0.131 |
| P7 | -0.328 | 0.048 | 0.014 | 0.954 | -0.052 | 0.771 |
| P3 | -0.462 | 0.005 * | -0.129 | 0.450 | -0.271 | 0.104 |
| Pz | -0.405 | 0.015 | -0.205 | 0.222 | -0.214 | 0.201 |
| P4 | -0.443 | 0.008 * | -0.109 | 0.523 | -0.148 | 0.383 |
| P8 | -0.129 | 0.450 | 0.052 | 0.771 | 0.129 | 0.450 |
| O1 | -0.129 | 0.450 | 0.157 | 0.352 | 0.186 | 0.270 |
| O2 | 0.024 | 0.907 | 0.224 | 0.181 | 0.367 | 0.027 |
| All electrodes | -0.290 | 0.081 | -0.033 | 0.862 | -0.100 | 0.561 |

**Table S1. Correlations between condition-related changes (MW – BF) in EEG complexity (HDF, LZC, and SampEn) and drowsiness level, for non-matched trial counts (non-MTC) approach.** Kendall's correlation was performed for each electrode between the difference in complexity and the drowsiness levels across subjects. Kendall's correlation was also applied for all electrodes between the difference in complexity (the average values across all electrodes) and the drowsiness levels across subjects. The $\tau$-values and the corresponding *p*-values (the original *p*-values, uncorrected by FDR) are reported. Asterisks indicate the presence of significance after FDR correction (threshold: 0.05). After FDR correction, electrodes C3, P3 and P4 show a negative correlation between the difference in HFD and the drowsiness level, with statistical significance.

| Electrode | HFD | | LZC | | SampEn | |
|---|---|---|---|---|---|---|
| | *τ* | *p* | *τ* | *p* | *τ* | *p* |
| Fp1 | -0.199 | 0.341 | 0.028 | 0.924 | 0.028 | 0.924 |
| Fp2 | -0.237 | 0.253 | 0.047 | 0.849 | 0.104 | 0.634 |
| F7 | -0.142 | 0.505 | 0.047 | 0.849 | 0.028 | 0.924 |
| F3 | -0.237 | 0.253 | -0.331 | 0.106 | -0.066 | 0.775 |
| Fz | -0.407 | 0.046 | -0.142 | 0.505 | -0.142 | 0.505 |
| F4 | -0.123 | 0.568 | -0.047 | 0.849 | 0.161 | 0.446 |
| F8 | -0.123 | 0.568 | 0.161 | 0.446 | 0.028 | 0.924 |
| T7 | -0.237 | 0.253 | -0.104 | 0.634 | 0.028 | 0.924 |
| C3 | -0.426 | 0.036 | -0.312 | 0.128 | -0.293 | 0.153 |
| Cz | -0.199 | 0.341 | -0.218 | 0.295 | -0.369 | 0.071 |
| C4 | -0.142 | 0.505 | -0.161 | 0.446 | -0.369 | 0.071 |
| T8 | -0.256 | 0.216 | -0.218 | 0.295 | -0.180 | 0.392 |
| P7 | -0.312 | 0.128 | -0.028 | 0.924 | 0.066 | 0.775 |
| P3 | -0.483 | 0.017 | -0.123 | 0.568 | -0.256 | 0.216 |
| Pz | -0.369 | 0.071 | -0.142 | 0.505 | -0.123 | 0.568 |
| P4 | -0.388 | 0.057 | -0.256 | 0.216 | -0.009 | 1.000 |
| P8 | -0.028 | 0.924 | 0.047 | 0.849 | 0.009 | 1.000 |
| O1 | -0.066 | 0.775 | 0.123 | 0.568 | 0.142 | 0.505 |
| O2 | 0.123 | 0.568 | 0.199 | 0.341 | 0.369 | 0.071 |
| All electrodes | -0.275 | 0.183 | 0.028 | 0.924 | -0.066 | 0.775 |

**Table S2. Correlations between condition-related changes (MW – BF) in EEG complexity (HDF, LZC, and SampEn) and drowsiness level, when adopting the matched trial counts (MTC) approach.** Kendall's correlation was performed for each electrode between the difference in complexity and the drowsiness levels across subjects. Kendall's correlation was also applied for all electrodes between the difference in complexity (the average values across all electrodes) and the drowsiness levels across subjects. The $\tau$-values and the corresponding *p*-values (the original *p*-values, uncorrected by FDR) are reported. After FDR correction (threshold: 0.05), no significant correlation was observed.

| one-sample *t*-test | HFD | LZC | SampEn |
|---|---|---|---|
| ***t*(24)** | -0.564 | 0.431 | -0.045 |
| ***p*** | 0.578 | 0.670 | 0.965 |
| **95% Confidence Interval** | [-0.143, 0.082] | [-0.093, 0.141] | [-0.126, 0.121] |

**Table S3. No significant correlation was observed between EEG complexity (HDF, LZC, and SampEn) and trial numbers.** Assuming that drowsiness increases over time, Spearman's correlation was analyzed between trial complexity and trial serial number per subject, and the *p(rho)*-value was obtained for each subject. Then a one-sample *t*-test was performed over the *p(rho)*-values and the results were reported.
